# Chronically-implanted Neuropixels probes enable high yield recordings in freely moving mice

**DOI:** 10.1101/406074

**Authors:** A.L. Juavinett, G. Bekheet, A.K. Churchland

## Abstract

The advent of high-yield electrophysiology using Neuropixels probes is now enabling researchers to simultaneously record hundreds of neurons with remarkably high signal to noise. However, these probes have not been comprehensively tested in freely moving mice. It is critical to study neural activity in unrestricted animals, and the field would benefit from the inclusion of ethological approaches to studying the neural circuitry of behavior. We therefore adapted Neuropixels probes for chronically-implanted experiments in freely moving mice. We demonstrate the ease and utility of this approach in recording hundreds of neurons across weeks, and provide the methodological details for other researchers to do the same. Importantly, our approach enables researchers to explant and reuse these valuable probes.

## Introduction

Observing behavior and recording neural activity in freely moving animals is crucial for our understanding of how the brain operates in the real world. Electrophysiology in freely moving rodents has been used to observe place and grid cell dynamics^1,2^, cortical dynamics during attentional control^3^, the role of oscillations during fear learning ^4^, whisking behavior during exploration ^5^, and more. Although freely moving recordings can be challenging, recording from unrestrained mice enables researchers to investigate behaviors that involve natural movements and offers ethologically-valid insight into neural activity^6^. Electrophysiology in freely moving animals is commonly performed with static electrode arrays or microdrives ^7–9^. These techniques have contributed much to the field, but are not at pace with the spatiotemporal coverage of cutting edge recording techniques, such as Neuropixels probes^10,11^.

### Neuropixels probes offer unprecedented insight into neural activity

Recent advancements in semiconductor technology have enabled the development of high-density silicone probes known as Neuropixels^10^. The linear recording shank can record from 384 contacts across 3.84 millimeters (selectable from 960 available sites on a 10 millimeter length shank). They have impressive signal-to-noise ratios (<8 μV RMS) as well as on-site amplification and digitization, enabling simultaneous recording of hundreds of cells across brain regions in an unprecedented low-noise, high-throughput manner. Importantly, methods have also been developed to automatically sort spikes from these recordings, and even correct for probe drift^12,13^.

Neuropixels probes have already proved invaluable for neuroscientists conducting acute experiments in mice, or chronic experiments in freely moving rats^10,14,15^. However, there is limited work with these probes in unrestrained mice^16^ although there is interest in behaviors and computations that involve movements of the animal’s head in space^15^. Further, although these probes have been very successful in freely moving rats^10,14^, there isn’t an established method to recover them after the experiment.

The opportunity to explant and reuse Neuropixels probes is essential because they are currently available only in very limited quantities, and when they are released, will be on the order of $1,000 each. Given the cost and very limited availability of the probes, many researchers will only be able to use them if it is possible to recycle them after experiments. We therefore sought to design an encasing for the Neuropixels probe that would allow experimenters to chronically implant it, run an experiment, and explant it for future experiments.

There are several considerations in designing a removable holder for chronic implants of Neuropixels probes in unrestrained mice. First, the current design of the probe has several components that need to be securely mounted onto the small mouse skull. Further, these sensitive onboard electronics need to be protected while the mouse is in its home cage. Most importantly, the shank of the probe must be secured to ensure consistent recordings across weeks of recording. In previous work, this required permanently mounting the biosensor using adhesive, which makes it nearly impossible to remove the probe afterwards^7^.

To address these needs, we designed the Apparatus to Mount Individual Electrodes (AMIE), an encasing that will protect the sensitive onboard electronics of the Neuropixels probe throughout long term, freely moving experiments. Moreover, the Neuropixels AMIE allows explantation and recycling. Our design and protocol is applicable to laboratories that wish to adapt the Neuropixels probe for recording in freely moving mice. Researchers that are using this technology in rats or acute mice may also find aspects of this approach useful.

With this design we have successfully recorded ∼100 neurons simultaneously from unrestrained mice while observing freely-moving behavior, and explanted the Neuropixels probe with a functioning recording shank.

## Results

### Design overview

The entire AMIE encasing weighs ∼1.5 g (with cement: ∼2.0 g) and is assembled from three parts: the Neuropixels probe, the internal mount (IM), and external casing (EC) (Figure 1a,b). The IM attaches directly to the Neuropixels PCB board with adhesive and is the core of the assembly (Figure 1a). On the backside of the IM is a slot for a stereotax adapter (SA) which allows for easy handling of the probe (Figure 1A). The IM attaches to the EC via a rail system (Figure 1b). During the implantation procedure, all adhesive binding the assembly to the rodent’s skull exclusively contacts the EC, which acts as a protective shell (Figure 1d).

**Figure 1.**
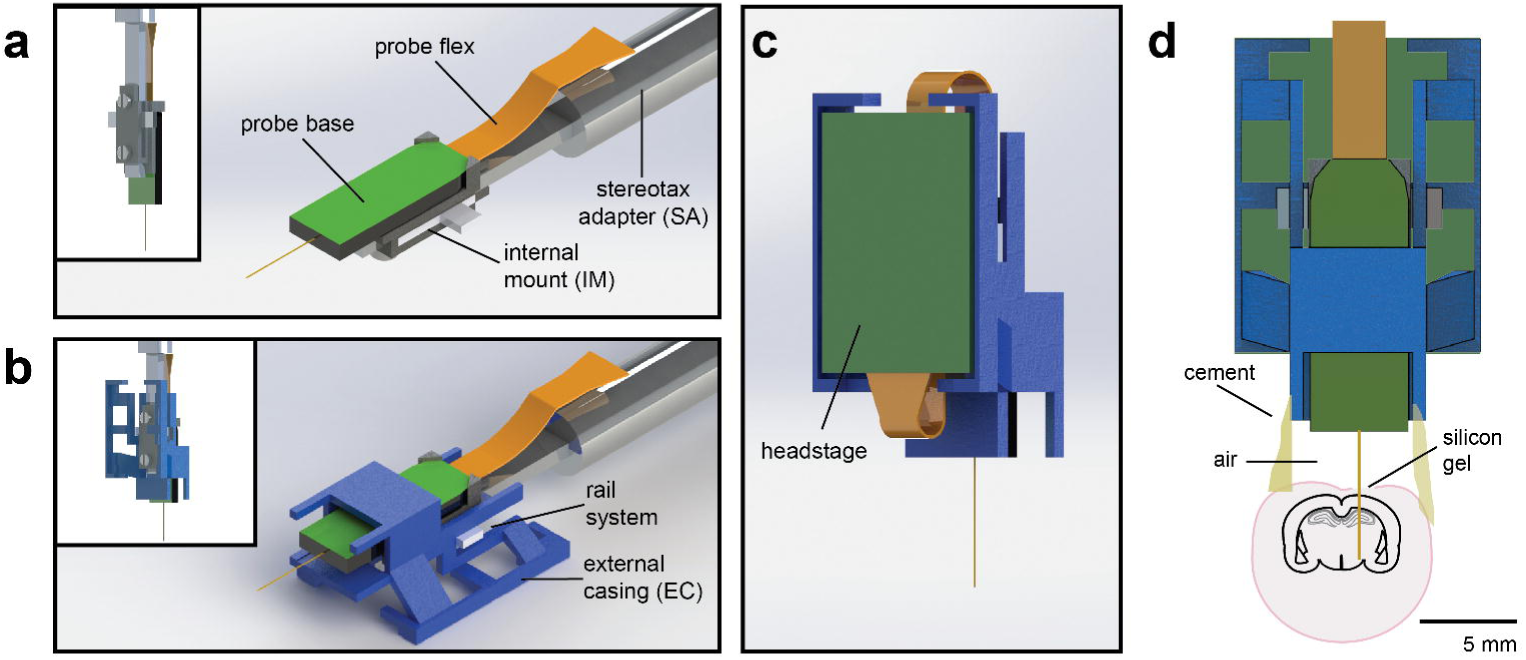
Schematic of Neuropixels AMIE. a. Probe base mounted onto 3D printed internal casing and attached to machined metal stereotax adapter *Inset:* Rear view, with screws that attach the internal mount (IM) to the stereotax adapter (SA). b. Entire assembly in a. within 3D printed external casing. *Inset:* Rear view c. The headstage is positioned on the back of the encasing, with the flex wrapped in an “S” shape. d. Entire assembly in relation to size of mouse brain and skull. The EC is attached to the skull with cement. Silicon gel is used to as an artificial dura to protect the open craniotomy

One difficulty in adapting the current Neuropixels design for freely moving experiments in mice is the ∼3 cm long flex cable attached to a 1 g headstage (see Jun et al. 2017 for details). In early testing, we suspended the flex and headstage above the mouse’s head during recording. However, we found that the flex very quickly twisted, potentially causing damage to it. In addition, the headstage added additional swinging weight above the mouse’s head. With these observations in mind, we designed the encasing with a space for the headstage to be semi-permanently affixed. The probe flex wraps in an “S” shape behind the implant, and attaches to the bottom (Figure 1c). In this way, the recording cable can be attached to the top of the implant, suspended above the mouse’s head.

Neuropixels 3A and 3B version probes were not designed for chronic implants in freely moving mice, and the entire probe assembly is quite bulky in comparison to a mouse’s head (Figure 1d, Jun et al. 2017). However, we have designed a very slim encasing for the probe, and have shown in our testing that mice adjust to the weight and size of the implant (Supplementary Video 1).

### Protocol overview

At least one day prior to implant, we attach the probe to the internal mount (Figure 2a). Silicon is added to further secure the base of the recording shank (Figure 2b). Once this is dry, the internal mount is slid into the rails of the external casing and secured with cement (Figure 2c,d). This cement will be drilled away in order to explant the probe. When the entire AMIE assembly is dry, it is ready to be implanted (Figure 2e). The surgery to implant the probe and encasing typically takes ∼3 hours (see Methods for details). During this surgery, a headbar can also be implanted, which does not interfere with the encasing. The external casing is the only part of the assembly that is attached to the skull (Figure 2f). In a typical experiment, we implant the probe and encasing without the headstage attached. We wait ∼3-4 days for the mouse to recover, and then add the headstage. The headstage can be removed after each experiment, if desired. After ∼1 day of habituation to the additional weight of the headstage (∼1 g), we begin recording during behavior.

**Figure 2.**
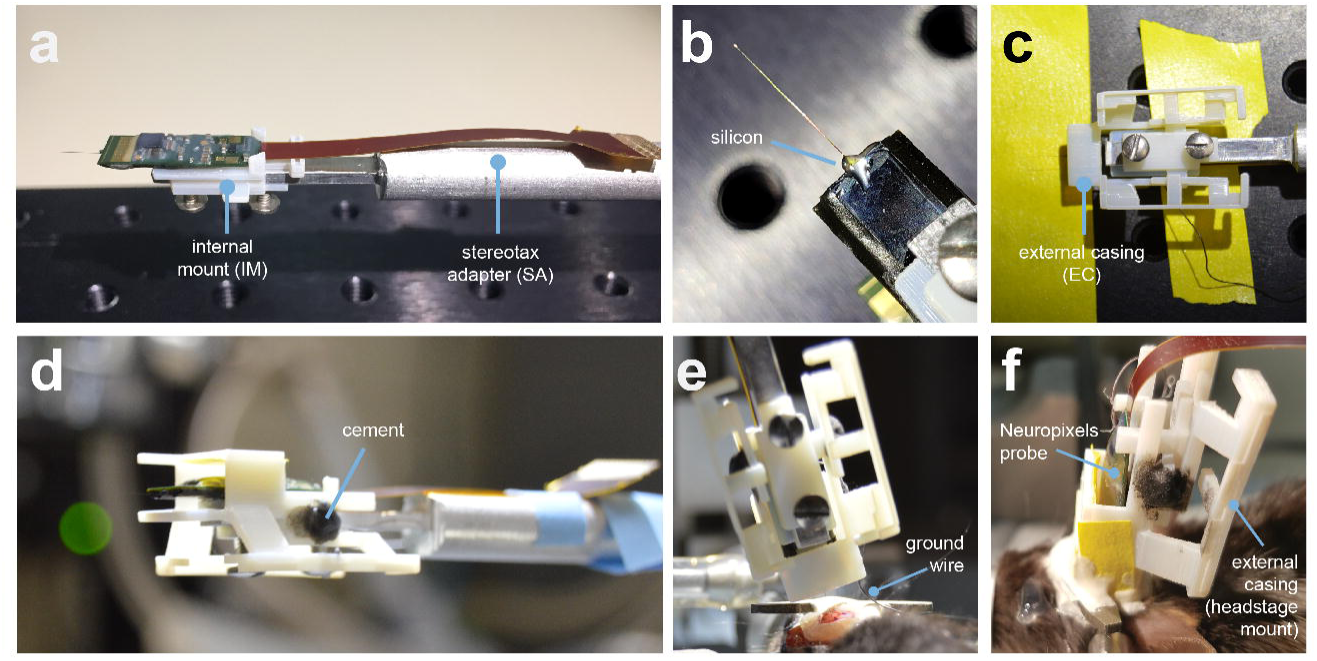
Mounting Neuropixels probe into the encasing. a. The internal mount (IM) is attached to the stereotax adapter with two screws, and probe is attached to the internal mount using an epoxy. **b.** The external case (EC) is attached to a breadboard, and the IM+probe assembly is carefully guided into the internal compartment of the EC (top view). c. Medical-grade silicon is added to the base of the shank to add extra support. d. After cementing the IM to the EC, the entire assembly is ready to be implanted. e. During surgery, the the shank is lowered into the brain (here at a ∼16 degree angle). The ground wire extends down the side of the implant and is attached to the ground screw. **f.** The entire encasing is attached to the headbar and skull using Metabond. Tape is added where necessary to add protec tion between the encasing and the skull. The stereotax adapter (not shown) is removed after this support structure is dry. The entire assembly is wrapped in Kapton tape after surgery.

### Mice are mobile with the implant

On the day after surgery, mice were already clearly mobile with the Neuropixels AMIE implant. Several days after surgery, we tested mice in an open arena to assess whether the implant impeded their behavior (Figure 4a,b)

**Figure 4.**
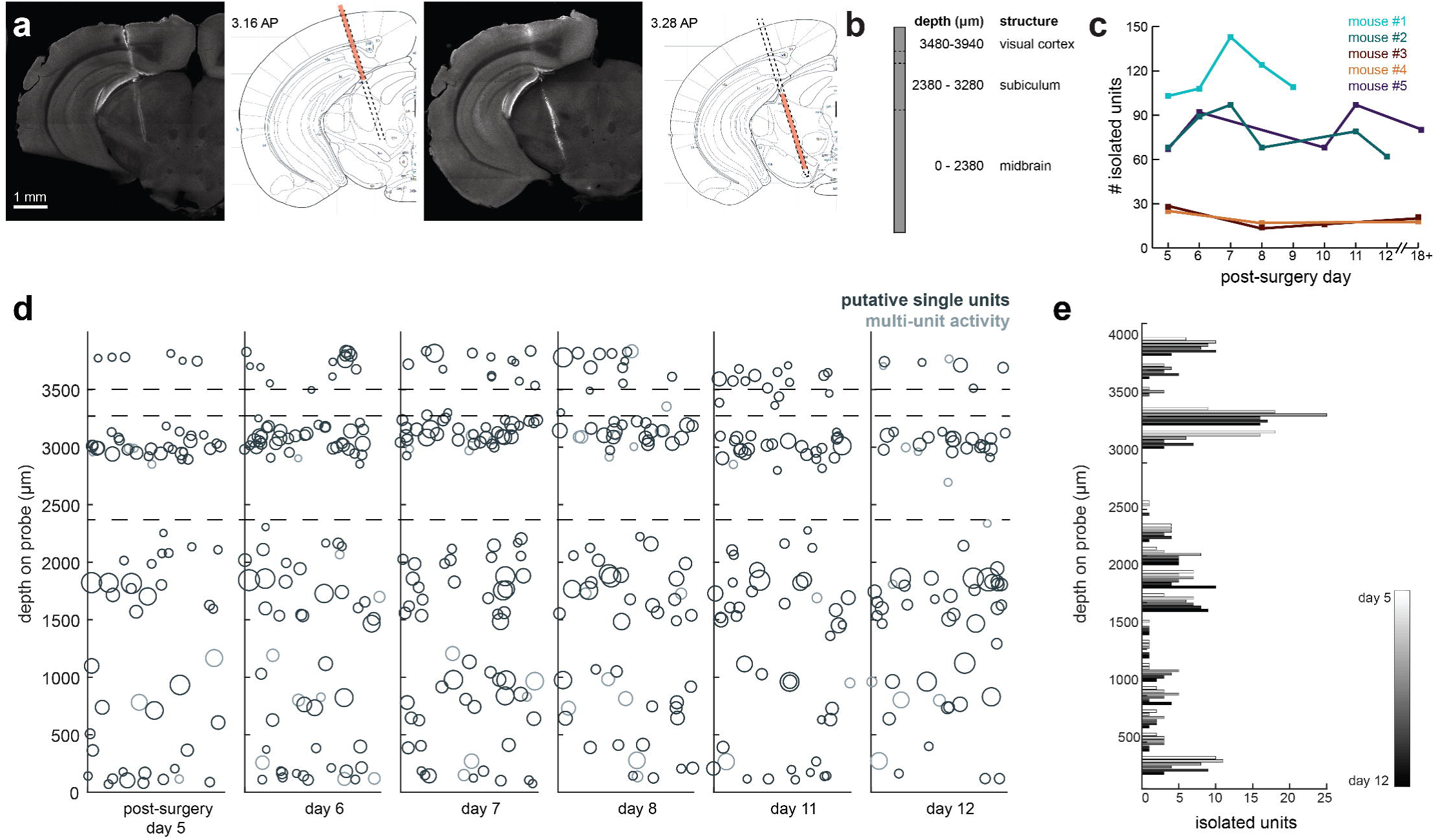
Chronic Neuropixel implants in cortex and subcortical regions can record -20-145 units across multiple days. a. Probe location, marked with Oil. Sections from Paxinos & Franklin atlas provided for reference. Mouse was implanted with a probe in visual cortex, hippocampus (subiculum), and the midbrain. **b.** Sche matic of probe depth with brain structures. **C.** Number of isolated units across recording days for 4 different mice. Mouse#1 &#2 were Option 1 probes (384 channels), Mouse#3 and#4 were Option 4 probes (276 channels). Mouse#2 is featured in the other panels of this figure. d. Scatter plot of of units across days for Mouse#2. Size of circles denotes number of waveforms assigned to that unit. X axis is random for visualization. e. Histogram of# of isolated units across days and across depth for Mouse#2.

Even while tethered, implanted mice are very agile (Supplementary Video 1). Implanted and naïve (unimplanted) mice spent similar percentages of their time moving around the rig (four mice, two 30-180 second samples per mouse; Figure 3d). The maximum velocities and acceleration in the open field of implanted mice were not statistically different than naive mice (4 mice, 2 samples per mouse; Supplementary Table 1; Figure 3d).

**Figure 3.**
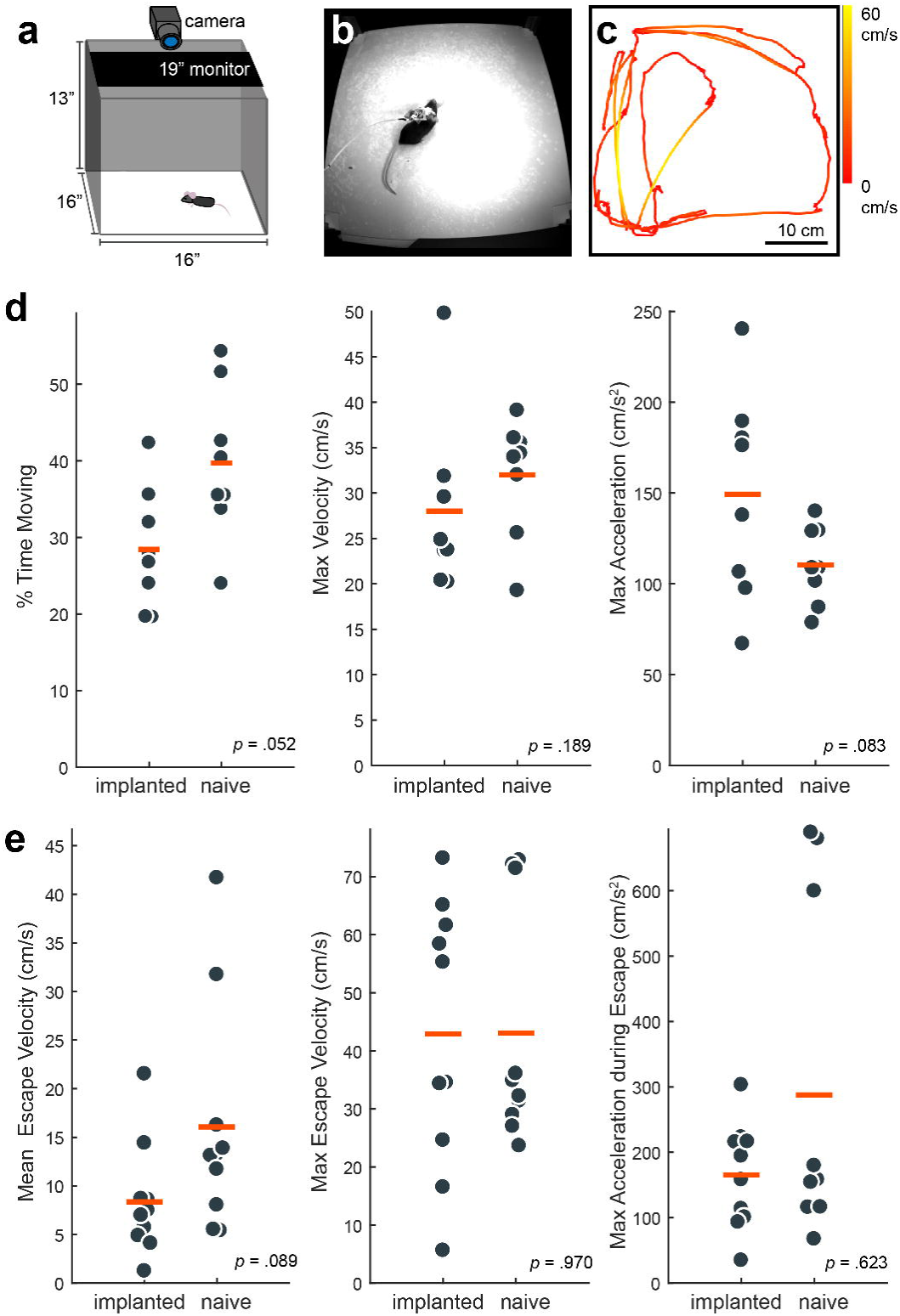
Behavior in implanted mice is comparable to naive mice. **a.** Behavioral testing arena, with a camera to track the position of the mouse and a monitor on top to present visual stimuli. **b.** Snapshot of mouse with implant in arena. **c.** Sample tracking of two minutes of open field behavior in an implanted mouse. Color of the line indicates the velocity of the mouse. **d.** Open field behav ior of implanted vs. naive mice. Random 30-180 second exerpts of behavior (N=B samples per group) in the open field were used to calculate a percent time moving(> 5 cm/s), max velocity, and max acceleration. Differences are consid ered significant at p <.0167 due to multiple comparisons. **e.** Visual-looming evoked behavior of implanted vs. naive mice. A dark dot of linearly increasing diameter(40 cm/s) was presented over the mouse’s head to evoke an escape response. The mean velocity, max velocity, and max acceleration during these responses is presented here. In both **d.** and **e.,** the orange line indicates the group mean. P values as computed by a two-sided Wilcoxon Rank Sum test are reported on each panel.

In addition, implanted and naive mice responded with comparable vigor to overhead visual looming stimuli (Figure 3e; Supplementary Table 2), which are known to elicit strong escape responses^16–18^. However, some naive mice achieved higher max acceleration during their escapes, at values unobserved in implanted mice (Figure 3e). This could be because of the slight obstruction introduced by the tether or because of the weight of the implant assembly.

### Chronic recording allows for 60-100 simultaneously recorded neurons, across weeks

We recorded spiking activity across multiple brain areas during freely moving behavior over the course of 1-2 weeks. Figure 4 illustrates an experiment with the probe implanted in medial visual cortex, subiculum, and midbrain. We isolated ∼60-100 units for each session in this experiment (Figure 4d&e). The number of single units we were able to isolate ranged across mice and experiments from ∼20-145, but these numbers were fairly consistent within each mouse across recording sessions (Figure 4c). This variability is likely dependent on the probe that was used (Option 4 probes used in mouse #3 and #4 had 276 rather than 384 recordable channels), recording noise, and brain region. The absolute number of isolated units depends on the quality of the sorting and the experimenter’s manual curation of Kilosort output, which does present challenging edge cases, and can be tricky with drift in the experiment (although alternative spike sorting software such as JRCLUST have been developed to address experiment drift^12^. Overall, these numbers are less than has been previously reported with acute experiments in mice^10^, possibly because of the chronic recording environment or inability to completely reduce noise. The longest we left a probe in was 41 days, without any noticeable decay in the signal.

### Researchers can also conduct headfixed recordings to further characterize neurons

A major limitation of many chronic implant designs is that they do not enable researchers to also implant a headbar to restrain the animal. We found the ability to do this critical for two reasons. First, it allowed us to easily restrain the mouse during experiments, e.g. to attach/replace the headstage or fix twisting in the tether. Second, it allowed us to present additional stimuli after the freely moving recording to further characterize the brain regions that we were recording from (Figure 5). This made it possible to connect the neural responses obtained during an unrestrained, ethological task with those obtained during more traditional sensory electrophysiology context. This opportunity could prove critical in bridging observations from these two very different contexts which are normally studied in separate laboratories.

**Figure 5.**
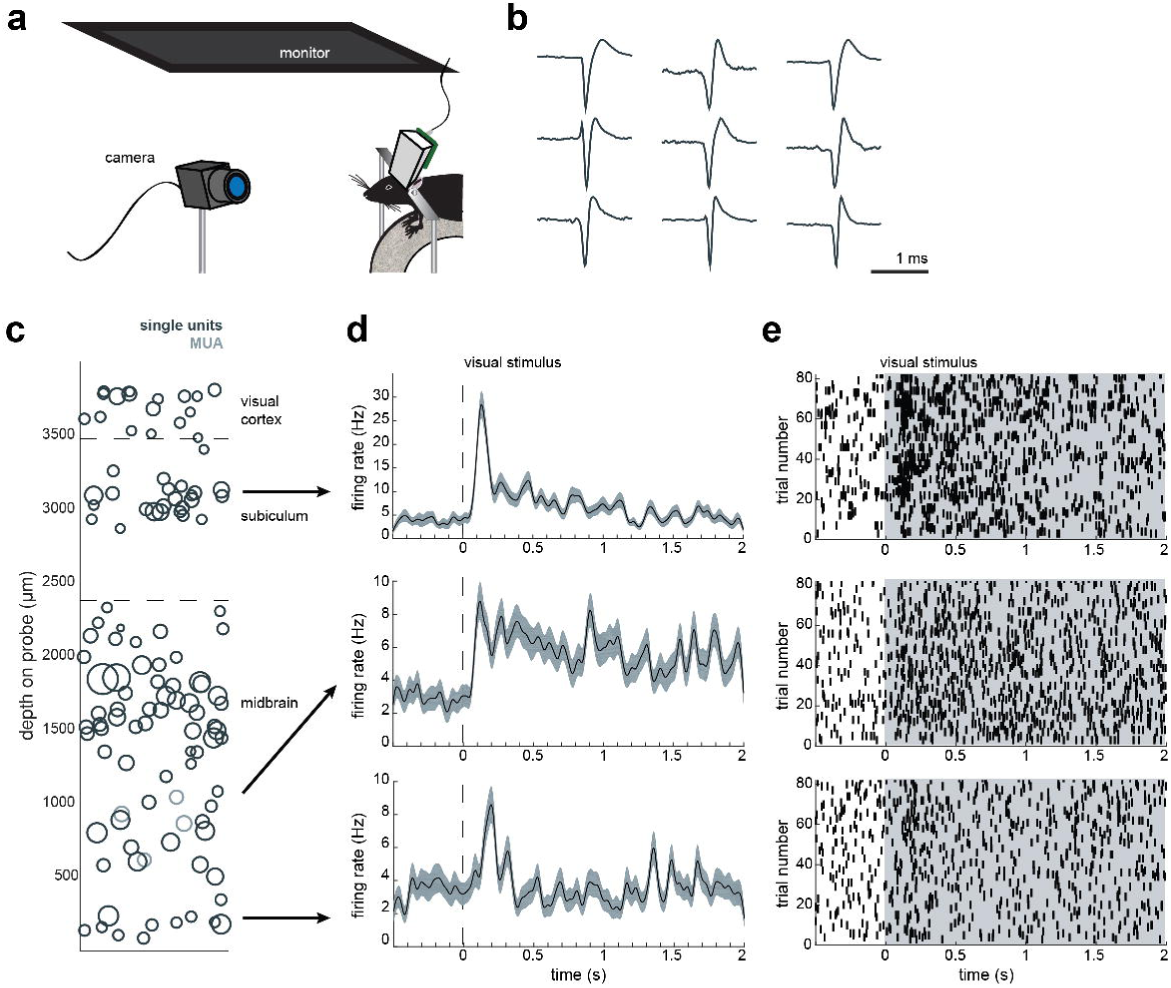
Implant design enables researchers to further characterize brain regions recorded during freely moving behavior. a. Schematic of headfixed setup. The mouse was implanted with a headbar (see Methods) enabling it to be restrained above a wheel/treadmill. Visual stimuli was presented on a monitor above the mouse’s head (similar to the unrestrained condition). The mouse’s pupil can be tracked with a high resolution IR camera, and movement can be tracked using a rotary encoder on a 3D printed wheel. b. Sample mean waveforms {n=500) from a headfixed recording, same mouse as Figure 4d (mouse #2). c. Distribution of sorted units across the probe, same mouse as Figure 4d (mouse #2). d. Peristimulus time histograms for three example neurons from different locations on the probe. The stimuli were a pseudorandomized set of 2-second full contrast sinusoidal drifting gratings in eight different directions. Shaded region is standard error of the mean. e. Raster plots for the neurons in d. Shaded area indicates the duration of the stimulus.

For example, after six days of recording freely moving behavior, we presented a battery of visual stimuli while the mouse was headfixed to determine whether cells were visually responsive (Figure 5a). We were able to isolate units in the restrained condition, just as in the freely moving condition (Figure 5b). The distribution of units was similar to previous experiments where the mouse was not restrained.

We also presented stimuli for retinotopic mapping and current source density analysis (to identify cortical layers; data not shown). Of course, researchers can present stimuli of their choice (visual or other) in the headfixed condition.

### Implant allows researchers to recover the probe after the experiment

Beyond providing a stable implant over many days, we also sought to design an implant that would allow for recycling of the Neuropixels probes. As demonstrated in Figure 1, the internal mount is separate from the external casing that is cemented to the mouse. After the completion of the experiment, researchers can drill away the cement and slowly remove the probe (see Methods and Figure 6a). This same probe, still attached to the internal mount, can then be re-secured within an external casing and implanted in another mouse.

**Figure 6.**
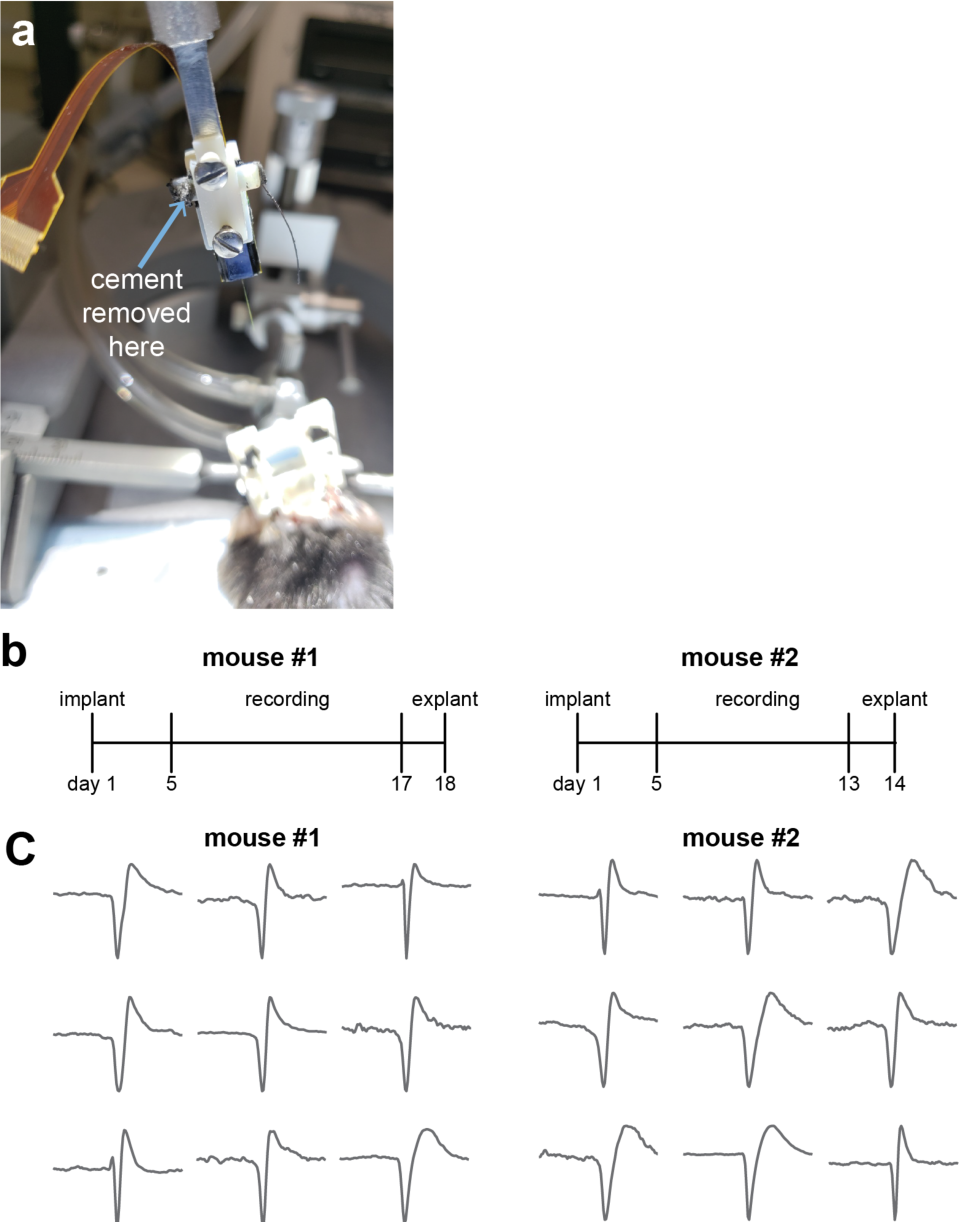
Implant design allows for probe explanation and subsequent re-implantation with the same probe. **a.** Example successful probe explantation. Cement is drilled away from the wings of the internal casing in order to remove the internal assembly from the external. **b**. Outline of experiment timing. The same probe was used in mouse #1 and mouse #2. **c**. Sample mean waveforms (n = 500) from each mouse. There was no noticeable change in signal quality in the second mouse.

In two mice with Option 3 Neuropixels probes (one shown in Figure 4), we were able to record from a mouse for over two weeks, explant the probe, and re-implant for a second experiment (Figure 6b). The quality of the recording did not noticeably change in the second mouse, and we were easily able to isolate clear units in both (Figure 6c).

Successful explant of probes depended on several factors. First, applying silicon to the base of the shank to add extra support appears to be necessary (Figure 1b). With silicon added to the base of the shank, 4/4 explant attempts were successful, whereas 1/6 explants were successful without the silicon (Table 1). Second, careful alignment of the probe, internal mount, and external casing will help ensure that the shank is being removed at the appropriate angle. Third, we only had success with Option 3 probes, suggesting that it may be easier with these. Fortunately, Option 3 probes are the version that will be on the market.

**Table 1.** Overview of experiments, with the Neuropixels probe option used and the outcome of the experiment. Starred mice are included in the paper.

| Mouse | Probe Option | Recordable Channels | Average # Isolated Units | Silicon on shank | Outcome |
| --- | --- | --- | --- | --- | --- |
| NP6* (Fig 4; Mouse #4) | 4 | 276 | 19.75 | no | Shank broke during explant |
| NP7* (Fig 4; Mouse #3) | 4 | 276 | 20 | no | Shank broke during freely moving recording |
| NP8* (Fig 4; Mouse #2) | 1 | 384 | 77.2 | no | Shank broke during explant |
| NP9* (Fig 4; Mouse #1) | 1 | 384 | 117.4 | no | Shank broke during explant |
| NP11 | 1 | 384 | - | no | Shank broke during freely moving recording |
| NP12 | 3 | 384 | - | no | Didn't recover from surgery, probe successfully explanted |
| NP13 | 3 | 384 |  | yes | Successful explant, re-implanted in NP15 |
| NP14* (Fig 6) | 3 | 384 | 80.6 | yes | Successful explant, re-implanted in NP16 |
| NP15 | 3 | 384 |  | yes | Successful explant |
| NP16* (Fig 6) | 3 | 384 |  | yes | Successful explant |

## Discussion

Here we present a significant advance in our ability to use and recycle high-density silicon probes such as Neuropixels. Our method allows researchers to perform recordings in both restrained and unrestrained conditions, and importantly, explant and reuse probes after experiments. Our hope is that this approach will enable researchers to capitalize on important technological advances to understand the complexity of brain activity during ethological behaviors.

Although Neuopixels probes were not designed for unrestrained recording in mice, we were able to adapt them for this purpose. We designed a slim enclosure for the probe as well as the headstage (Figure 1 &2), that mice can easily handle (Figure 3).

Unlike other electrophysiology systems, the current Neuropixels recording tether is not easily commutated due to heavy data demands (though attempts are underway). While this has not been a problem for recording from chronically-implanted rats in large arenas^10^, it can be challenging for recordings from mice in smaller arenas, requiring constant monitoring of the mouse’s position and occasional intervention from the experimenter to untangle the cord. In our experience, this has been manageable, and here we report similar behavior in implanted and naive mice (Figure 3).

The Neuropixels AMIE can be used to record in both restrained and unrestrained conditions, with similar yields in numbers of isolated units (Figures 4,5). The ability to restrain the mouse for passive stimulation enables researchers to obtain additional information about their recordings that may ultimately aid in uncovering the function of cells and brain regions. Remarkably, during our headfixed experiments we found that even cells deep in the midbrain showed clear visual responses to drifting gratings (Figure 5d,e). This demonstrates the power of Neuropixels to uncover signals for information in uncharted brain territories.

## Methods

### Printing and machining parts

To conduct this experiment, researchers will need Neuropixels probes. We recommend performing the entire process of preparing and implanting the probe using a dummy probe for practice. We printed and tested in VeroWhite material using a Stratasys Eden 260VS PolyJet 3D Printer with 16 μm resolution. The stereotax adaptor should be machined from aluminum or stainless steel. All designs can be found on the CSHL repository (http://repository.cshl.edu/36808/) as well as on Github (https://github.com/churchlandlab/ChronicNeuropixels).

### Mounting the probe

First, the internal mount is secured to the stereotax adapter (SA) using two 2-56A screws (Amazon, B00F34U238). As depicted in Figure 2a, we then attached the Neuropixels probe to the internal mount (IM) using Loctite Instant Adhesive 495 (ULINE S-17190). Using a needle, we applied a medical-grade clear silicon adhesive, Mastersil 912MED, to the base of the shank (Figure 2b). The IM & probe was slid into the rails of the external casing (EC), and secured with cement (Figure 2c&d).

### Surgical methods

All surgical and behavioral procedures conformed to the 316 guidelines established by the National Institutes of Health and were approved by the Institutional 317 Animal Care and Use Committee of Cold Spring Harbor Laboratory. We used male 3-4 month old C57/BL6 mice (Jackson Laboratories, 000664). Male mice were used because they are typically larger, and we expected that they would better handle the weight of the implant. Mice were given medicated (carprofen) food cups (MediGel CPF, Clear H20 74-05-5022) 1-2 days prior to surgery.

During surgery, the mouse was anesthetized with isoflurane. We cut away the skin and cleared any connective tissue. Tissue at the edges of the skull was glued down with Vetbond (Santa Cruz Biotechnology, cat. no. sc-361931). The skull was cleared and dried, using a skull scraper or blade to add additional texture. A boomerang shaped custom Titanium headbar was cemented to the skull, just posterior to the eyes, near Bregma. A burr hole was drilled for the ground screw, which was carefully screwed into the skull. We applied Optibond Solo Plus (Kerr, cat No. 31514) to the skull, and used UV light to cure it. We used Charisma (Net32, cat. No. 66000085) to create a base for the implant, and add additional support around the ground screw. Using a dental drill, a small craniotomy (1-2 mm) was made over visual cortex (2-2.5 ML,-3.4-3.5 AP relative to Bregma). The entire Neuropixels assembly (SA, IM, and EC) was placed in the stereotax and the shank was slowly lowered into the brain at a ∼16 degree angle (Figure 2e). The ground wire is wrapped around the ground screw, and Metabond cement was carefully applied to attach the EC to the skull. The entire assembly was wrapped in Kapton Tape (ULINE S-7595) and the mouse was allowed to recover for 3-4 days.

Once the mouse recovered, we removed the tape and added the headstage to the back of the implant. The entire assembly was re-wrapped with tape. On the next day, we began behavioral testing.

### Behavioral data

To compare the behavior of implanted mice with naive/unimplanted mice, we tracked mice using a Basler Pylon camera and Ethovision XT13 in a 16” × 16” open arena. For open field tests, unimplanted mice were allowed to explore a bare arena for 15 minutes. Implanted mice were tested in an arena with an inset nest; the data presented here are random excerpts of the mouse’s activity while outside of the nest. We excerpted the same length time segments (4 mice, 2 samples each) from the unimplanted mice for comparison.

We employed a Kolmogorov-Smirnov test to determine that the values for each of the open field and visual looming behavioral metrics were not normally distributed. Therefore, we used a two-sided Wilcoxon rank sum test in order to test differences between the implanted and unimplanted groups (Supplementary Tables 1&2).

### Visual stimulation

For visually-evoked responses during freely moving behavior (Figure 4), a linearly expanding dot (40 cm/s) was presented on a monitor directly over the mouse’s head. This stimulus is known to elicit an escape response in mice ^17,18^. Unimplanted mice could escape into a small nest: a triangular prism with a 13 cm opening. Implanted mice could escape into a nest inset into the wall. For visually-evoked responses during head restraint (Figure 6), a set of full contrast, full field drifting gratings in eight different directions (10 repeats) were presented above the mouse’s head while the mouse was free to move on a wheel.

### Electrophysiology data

Electrophysiology data was collected with SpikeGLX (Bill Karsh, https://github.com/billkarsh/SpikeGLX). The data were first median subtracted across channels and time (see Jun et al., 2017b) and then sorted with Kilosort spike sorting software ^13^ and manually curated using phy (https://github.com/kwikteam/phy). Additional analyses and plotting with data were done with MATLAB code modified from N. Steinmetz (https://github.com/cortex-lab/spikes).

### Probe explantation

To explant the probe, we first anesthetized the mouse with isoflurane and loosely positioned the mouse into the earbars. The SA was placed in the stereotax and aligned with its slot in the IM. We carefully lowered the SA into the IM, and put the two screws back into place. It was important that the SA was properly aligned with the IM so that no unnecessary tension was placed on the implant. We carefully drilled away the cement at the boundary of the IM and EC, unraveled or cut the ground wire, and slowly raised the SA+IM+probe assembly. The mouse was perfused and the brain was fixed in 4% PFA for sectioning.

### Protocol, code, and data availability

A detailed surgical protocol for mounting, implanting, and explanating the probe will be uploaded to the Nature Protocols Exchange. Code to generate the figures here will be provided on Github. Behavioral and electrophysiological data will be stored on a dedicated repository that is maintained by CSHL. Files will be linked from a lab webpage that is used exclusively for this purpose: http://churchlandlab.labsites.cshl.edu/code.

## Acknowledgements

This work represents the collective input and knowledge of a burgeoning Neuropixels community.

We would like to acknowledge Tim Harris for his leadership on the development of the Neuropixels probes and his constant encouragement of this project.

The UCL Neuropixels course, taught by Nick Steinmetz, Matteo Carandini, Andrew Peters, Adam Kampff, was imperative in getting this project off the ground (http://www.ucl.ac.uk/neuropixels/courses). In particular, we would like to thank Nick Steinmetz for his critically important feedback, code, and upkeep of the Neuropixels Github page (https://neuropix.cortexlab.net).

We would also like to acknowledge Claudia Boehm and Albert Lee (Janelia Research Campus) for allowing us to observe their rat Neuropixels implant. Their protocol served as an important starting point for the protocol we developed in mice.

We have also benefitted from troubleshooting help from many individuals, including Wade Sun, James Jun, Marius Bauza, and Bill Karsh (SpikeGLX).

We are also grateful to the CSHL Undergraduate Research Program, which yearly provides a diverse group of students with funding and resources to complete invaluable research experiences at CSHL. This program funded G.B. for his initial summer in our lab.

We welcome feedback from the community regarding the diversity of methods used to implant and record with these probes.

## Supplementary Material

**Table S1.** Statistics for open field behavior. *N* = 8 for both groups. SD is standard deviation, *Z* statistics and *p* values are calculated from two-sided Wilcoxon Rank Sum, r is measure of effect size, calculated by 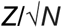.

|  |  | mean | SD | Z | p | r |
| --- | --- | --- | --- | --- | --- | --- |
| % moving | implanted | 28.45 | 7.88 | -1.94 | 0.052 | -0.686 |
|  | naive | 39.7 | 9.86 |  |  |  |
| max velocity | implanted | 28.3 | 9.46 | -1.31 | 0.189 | -0.463 |
|  | naive | 30.86 | 5.57 |  |  |  |
| max acceleration | implanted | 154.41 | 54.69 | 1.73 | 0.083 | 0.612 |
|  | naive | 106.23 | 27.42 |  |  |  |

**Table S2.** Statistics for visual looming evoked responses. N = 10 for both groups. SD is standard deviation, *Z* statistics and *p* values are calculated from two-sided Wilcoxon Rank Sum, r is measure of effect size, calculated by 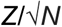.

|  |  | mean | SD | Z | p | r |
| --- | --- | --- | --- | --- | --- | --- |
| mean velocity | implanted | 8.37 | 5.78 | -1.70 | 0.089 | -0.538 |
|  | naive | 16.06 | 11.70 |  |  |  |
| max velocity | implanted | 42.93 | 22.89 | -0.038 | 0.970 | -0.012 |
|  | naive | 43.06 | 20.38 |  |  |  |
| max acceleration | implanted | 165.39 | 79.92 | -0.491 | 0.623 | -0.155 |
|  | naive | 287.78 | 257.10 |  |  |  |

